# Detecting tandem repeat expansions in cohorts sequenced with short-read sequencing data

**DOI:** 10.1101/157792

**Authors:** Rick M Tankard, Mark F Bennett, Peter Degorski, Martin B Delatycki, Paul J Lockhart, Melanie Bahlo

## Abstract

Repeat expansions cause over 30, predominantly neurogenetic, inherited disorders. These can present with overlapping clinical phenotypes, making molecular diagnosis challenging. Single gene or small panel PCR-based methods are employed to identify the precise genetic cause, but can be slow and costly, and often yield no result. Genomic analysis via whole exome and whole genome sequencing (WES and WGS) is being increasingly performed to diagnose genetic disorders. However, until recently analysis protocols could not identify repeat expansions in these datasets.

A new method, called exSTRa (**e**xpanded **S**hort **T**andem **R**epeat **a**lgorithm) for the identification of repeat expansions using either WES or WGS was developed and performance of exSTRa was assessed in a simulation study. In addition, four retrospective cohorts of individuals with eleven different known repeat expansion disorders were analysed with the new method. Results were assessed by comparing to known disease status. Performance was also compared to three other analysis methods (ExpansionHunter, STRetch and TREDPARSE), which were developed specifically for WGS data. Expansions in the STR loci assessed were successfully identified in WES and WGS datasets by all four methods, with high specificity and sensitivity, excepting the FRAXA STR where expansions were unlikely to be detected. Overall exSTRa demonstrated more robust/superior performance for WES data in comparison to the other three methods. exSTRa can be applied to existing WES or WGS data to identify likely repeat expansions and can be used to investigate any STR of interest, by specifying location and repeat motif. We demonstrate that methods such as exSTRa can be effectively utilized as a screening tool to interrogate WES data generated with PCR-based library preparations and WGS data generated using either PCR-based or PCR-free library protocols, for repeat expansions which can then be followed up with specific diagnostic tests. exSTRa is available via GitHub (https://github.com/bahlolab/exSTRa).

## Introduction

Thousands of short tandem repeats (STRs), also called microsatellites, are scattered throughout the human genome. STRs vary in size but are commonly defined as having a repeat motif 2-6 base pairs (bps) in size. They are underrepresented in the coding regions of the human genome^1^, despite the vast majority being population polymorphisms with no, or very little, phenotypic consequence. STRs were used as genetic markers for linkage mapping for human studies for many years, and continue to be used, but primarily for non-human studies. A subset of STRs can however cause disease. Pathogenic STRs have either one or two alleles, depending on the genetic model, that exceed some threshold for biological tolerance. These diseases are known as repeat expansion disorders. The abnormal STR allele(s), may affect gene expression levels, cause premature truncation of the protein or result in aberrant protein folding.^2^ Repeat expansions at different STR loci share biological consequences. Common disease mechanisms mediated by repeat expansion disorders include Repeat-associated non-AUG translation and MBNL spliceosome interference, for example caused by CUG expansions in Myotonic Dystrophy Type 1 (DM1). These mechanisms are reviewed in Hannan^3^.

Repeat expansions cause ~30 inherited germline human disorders, predominantly neurogenetic diseases most often presenting with ataxia as a clinical feature. The size of pathogenic allele varies from ~60 repeats observed in the gene encoding the Calcium Voltage-Gated Channel Subunit Alpha1 A (*CACNA1A*) to several thousand repeats observed in the gene encoding the Calcium Voltage-Gated Channel Subunit Alpha1 A (*C9orf72*) (Table 1). Remarkably 12 repeat expansions have now been identified as causing dominant forms of spinocerebellar ataxias. Other disorders caused by repeat expansions include fragile X syndrome (OMIM #300624, a repeat in the 5’UTR of *FMR1*), Huntington Disease (OMIM #606438, a repeat in exon 1 of *HTT*), myotonic dystrophy (OMIM #602668, repeats in *DMPK* and *ZNF9*), fronto-temporal dementia and amyotrophic lateral sclerosis 1 (OMIM #105550, a 6-mer repeat in *C9orf72*) and Unverricht-Lundborg disease, a severe myoclonic epilepsy (OMIM #254800, in *CSTB*). The genetic mode of inheritance encompasses autosomal dominant (e.g. SCA1, OMIM #164400) and recessive (e.g. Freidreich ataxia, OMIM #229300), as well as X-linked recessive (e.g. fragile X syndrome, OMIM #300624). Novel pathogenic alleles underlying repeat expansion disorders continue to be discovered, with the two most recently described STRs being pentamer repeats^4; 5^. A selected list of repeat expansion disorders are shown in Table 1.

**Table 1.**
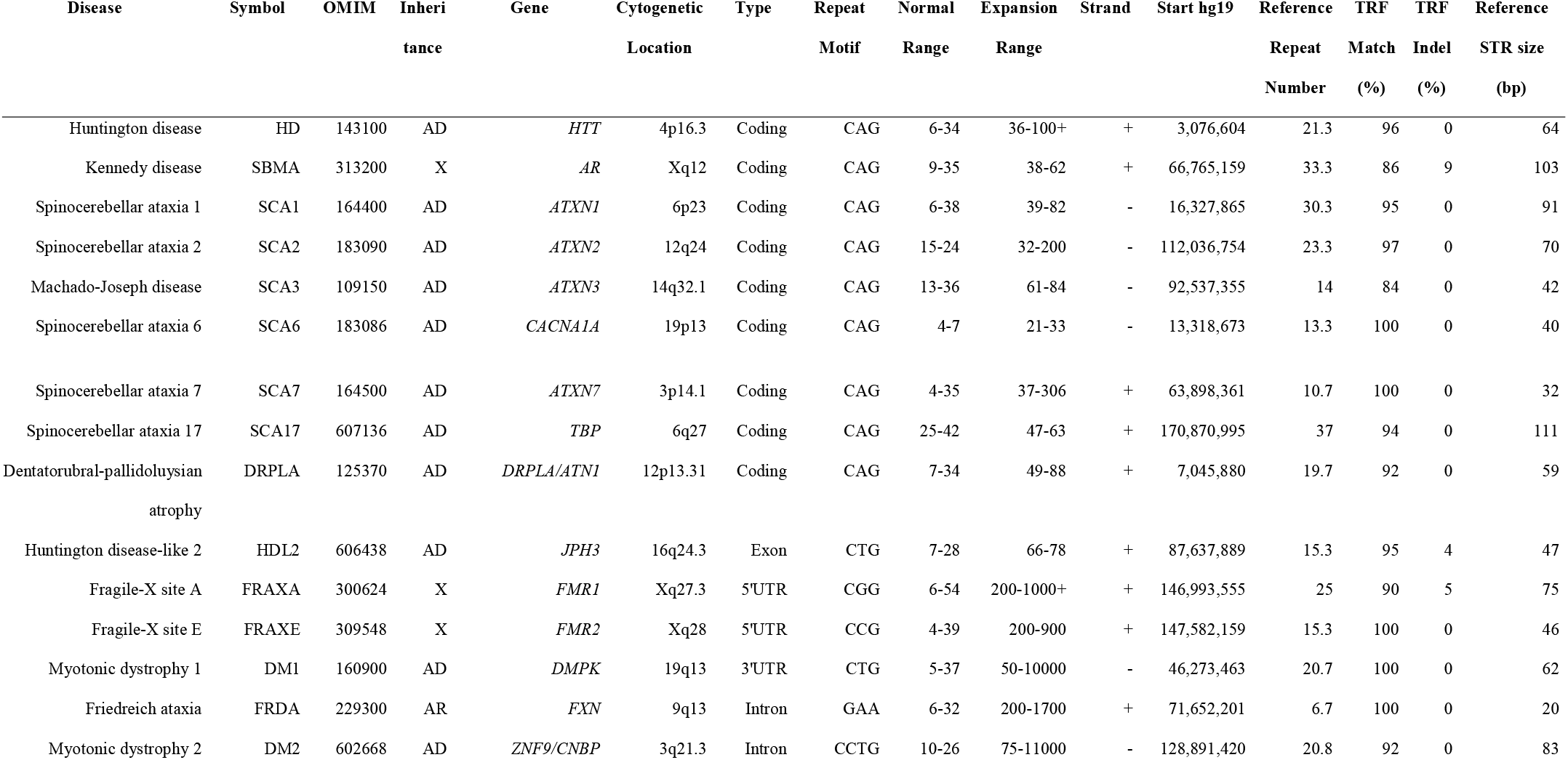
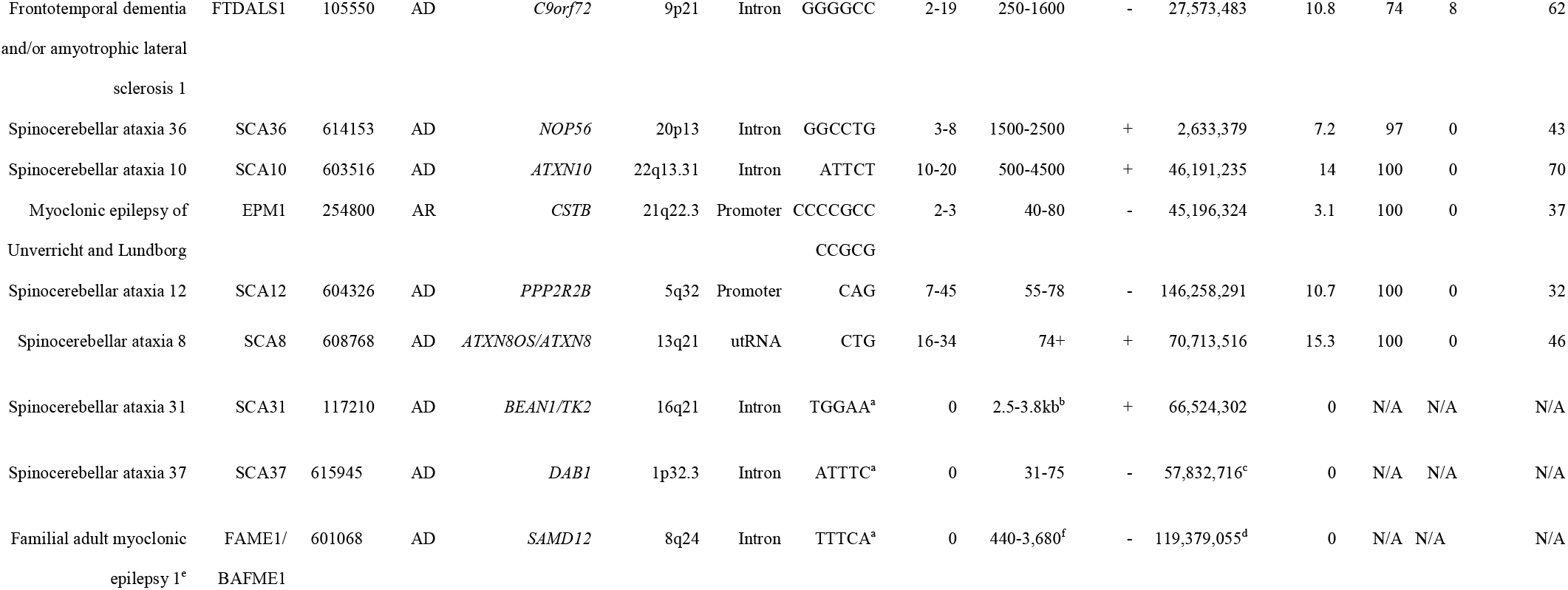
Short tandem repeat loci information for STRs causing neurogenetic disorders. TRF, Tandem Repeats Finder (Benson et al, 1999). TRF match and TRF indel describe the purity of the repeat. AD, autosomal dominant; X, X-linked; AR, autosomal recessive; UTR, untranslated region. ^a^These repeat expansions are novel insertions and thus not repesented in the reference genome at their respective locations. ^b^SCA31 is caused by the insertion of a complex repeat containing (TGGAA)_n_;; hence the length is given in as the length of the expanded repeats in bps, instead of repeat number. ^c^The SCA37 physical map location is given at the reference (ATTTT)n repeat, where affected individuals have the pathogenic (ATTTC)_n_ insered. ^d^The FAME1 physical map location is given as the position of the reference (TTTTA)_n_ repeat, at which affected individuals have (TTTCA)_n_ inserted, ^e^lshiura et al. identified similar expansions assocatioaed with FAME6 and FAME7, in the genes TNRC6A and RAPGEF2 respectively, but only in single families. These have not been listed. ^f^The FAME1 repeat size is the estimated size of the combined expanded (TTTCA)_n_ and the (TTTTA)_n_ reference repeat.

Many repeat expansion disorders show anticipation; a phenomenon whereby younger generations are affected by earlier age of onset. Anticipation is usually caused by an increase in repeat size between generations. When anticipation is observed it indicates that a search for repeat expansions as the cause of disease is warranted.

Friedreich ataxia is the most common of the recessive repeat expansion disorders, with a disease prevalence of 3 to 4/100,000 but with a carrier frequency of 1/100.^6^ Fragile X syndrome is the most common cause of inherited intellectual disability and affects ~1/5000 individuals.^7; 8^ Hence these diseases as a whole contribute significantly to the overall Mendelian disease burden in human populations.

Diagnostic identification of repeat expansions can be time consuming and costly. Current medical diagnosis consists of precise PCR or Southern blot assay, which require diagnostic laboratories that have refined these assays for each different repeat expansion. The clinician has to determine which repeat expansions are most likely to be relevant and submit the patient’s DNA to appropriate laboratories. This can be difficult, given the phenotypic overlap between the different STRs, the potential heterogeneity in the symptoms and the variation in penetrance and age of onset, which is also dependent on the size of the allele and effect of modifier genes.^9; 10^ In addition, up to 50% of individuals with a diagnosis of ataxia may be due to other mutation types, such as single nucleotide variants (SNVs) and short insertion/deletions (indels).^11^ Therefore, molecular diagnosis of these disorders often also requires conventional sequencing of candidate genes, either by Sanger, targeted panel or Next Generation Sequencing (NGS) methods.

Short-read NGS data, such as that generated by the Illumina sequencing platform, is currently predominant in both research and clinical diagnostic applications. Moreover, Whole Genome Sequencing (WGS) is now an affordable technology, gradually replacing whole exome sequencing (WES) for clinical genomics. Illumina’s HiSeq X and NovaSeq platforms are currently the most commonly used platform for the generation of human WGS data and in particular clinical human genome sequencing with low error rates and well-documented, consistent, performance.

Illumina HiSeq X data reads are 150 bp in length and are designed so that the reads are transcribed facing each other, where the template DNA predominantly has a small gap between the reads that is not sequenced. This gap can vary in size, but standard library preparation methodologies generate insert fragment lengths of ~350 bps, resulting in a gap of ~50bp.

Standard clinical diagnostic pipelines focus on the identification of SNVs and indels. Bioinformatic tools have been developed to genotype STRs, but are almost entirely confined to those STR alleles that are spanned by reads.^12–16^ Pathogenic repeat expansions are usually significantly longer than the reads generated by short-read sequencing platforms such as Illumina, and may be longer than the library insert fragments lengths. Therefore, the short reads cannot span many pathogenic repeat expansion alleles, such as those that cause SCA2 (OMIM #183090), or SCA7 (OMIM #164500, Table 1). Furthermore some of these reads are not mapped, or poorly mapped, to the STR allele, due to sequencing bias and alignment issues such as: (i) the repetitive nature of the repeat itself where the expanded alleles require alignments of additional repetitive bases, (ii) multiple occurrences of the same repeat throughout the genome, leading to multi-mapping reads, and (iii) GC bias. Despite this, these data do still carry information about the expanded allele with a larger number of reads mapping to the STR for an expanded allele than expected, based on the reference STR allele lengths.

Several methods now describe the detection of repeat expansion in short read NGS data. These include ExpansionHunter^17^, STRetch^18^ and TREDPARSE^19^, reviewed in Bahlo et al^20^. These methods are focused on detection of repeat expansions in whole genome sequencing data, with a preference for PCR-free library free protocols. ExpansionHunter and TREDPARSE determine whether an individual has an expansion based on pre-determined thresholds, however TREDPARSE also has a likelihood ratio test with a likelihood framework that determines the genetic model and the likelihood of expansion. STRetch uses a genome reference augmented with decoy chromosomes, consisting of long stretches of all 1 to 6 bp repeat expansions to competitively attract long repeats. None of these methods have been assessed for performance in comparison to each other or in WES data.

Here we describe the development of the STR repeat expansion-calling algorithm, exSTRa (**ex**panded **STR a**lgorithm), which detects expanded repeat expansion allele(s) at repeat expansion loci, specified by the user, in cohorts of sequenced individuals. We demonstrate the utility of the method with twelve different verified repeat expansion disorders. exSTRa is designed to be applied to cohorts of individuals without requiring a set of controls. This is because exSTRa is designed as an outlier detection test, where the majority of individuals (>85%) are assumed to have normal length alleles at a particular repeat expansion locus. This assumption is robust for the majority of disease cohorts, even spinocerebellar ataxias. exSTRa also generates unique empirical cumulative distribution function (ECDF) plots of individual’s repeat motif distributions, plotted for all individuals in a cohort, facilitating QC for batch effects and validity of assumptions. We demonstrate for the first time, that repeat expansion detection is possible with WES data and further demonstrate on additional STR loci, that PCR-based library preparation WGS, whilst inferior to PCR-free library preparation WGS data, can be used to confidently interrogate most known STR loci. This will enable researchers to interrogate the thousands of existing NGS datasets for repeat expansions at known repeat loci or any other loci they wish to investigate.

## Methods

### Study cohorts and next-generation sequencing data generation

Individuals with already diagnosed repeat expansion disorders were recruited for this study. The repeat expansion status was verified via standard diagnostic STR-specific PCR-based assays. Individuals affected by neurogenetics disorders not due to known repeat expansions were recruited as controls. These individuals were not tested for any of the known repeat expansion loci with standard methods as none of them are affected by symptoms that are typical of expansion disorders such as ataxia. All individuals were recruited at the Murdoch Children’s Research Institute, and provided written informed consent (Human Research Ethics Committee #28097, #25043 and #22073).

Four cohorts underwent different types of NGS, with some individuals being sequenced multiple times. Individuals were sequenced with either: (i) WES with the Agilent V5+UTR capture platform (4 repeat expansion patients, with 4 different expansion disorders, 58 controls), (ii) WGS with the TruSeq Nano protocol, which includes a PCR step to increase sequencing material (17 repeat expansion patients, with 8 different expansion disorders, 16 controls), or (iii) WGS with the PCR-free cohort consisting 118 individuals (52 females and 66 males). Samples in this cohort were either affected with the repeat expansion disorder, or carriers, for one of: FRAXA (15 expanded, 19 intermediate), FRDA (25), DM1 (17), HD (13), SCA1 (3), DRPLA (2), SBMA (1) and SCA3 (1), or relatives with no known expansion (22), with all samples sourced from the Coriell resource. The WES cohort is designated as WES_PCR. Two different cohorts were sequenced with protocol (ii). These are designated as WGS_PCR_1 and WGS_PCR_2. The WGS cohort was designated as WGS_PF. These cohorts are described in Table 2.

**Table 2.** Repeat type, genetic model, diseases, sample names and which cohorts samples appear in. Allele sizes are derived from standard laboratory tests for repeat expansions. Some individuals were not tested (Not sized) or the data was not available (not recorded). MOI = mode of inheritance (AD = autosomal dominant, X = X-linked, AR = autosomal recessive). Only the total number of controls are given denoted by (controls).

| 932 | Class | MOI | Diagnosis | Allele sizes | Gender | WES_PCR | WGS_PCR_1 | WGS_PCR_2 |
| --- | --- | --- | --- | --- | --- | --- | --- | --- |
| 933 | PolyQ | AD | HD | Not recorded | male | rptWEHI3 | HD-1 |  |
| 934 | PolyQ | AD | HD | 17,39 | female |  |  | WGSrpt_10 |
| 935 | PolyQ | AD | HD | 20,42 | male |  |  | WGSrpt_12 |
| 936 | PolyQ | AD | SCA1 | 36,52 | female | rptWEHI4 |  | WGSrpt_14 |
| 937 | PolyQ | AD | SCA1 | 30,45 | male |  |  | WGSrpt_16 |
| 938 | PolyQ | AD | SCA2 | 21,42 | female | rptWEHI1 | SCA2-1 | WGSrpt_18 |
| 939 | PolyQ | AD | SCA2 | 23,39 | male |  |  | WGSrpt_20 |
| 940 | PolyQ | AD | SCA6 | 11,22 | female | rptWEHI2 | SCA6-1 | WGSrpt_05 |
| 941 | PolyQ | AD | SCA6 | 10,21 | female |  |  | WGSrpt_07 |
| 942 | PolyQ | AD | SCA7 | 13,39 | female |  |  | WGSrpt_08 |
| 943 | 5'UTR | X | FRAXA | Not sized | male |  |  | WGSrpt_17 |
| 944 | 5'UTR | X | FRAXA | 613-1680 | male |  |  | WGSrpt_19 |
| 945 | 5'UTR | X | FRAXA (pre) | ~100 | female |  |  | WGSrpt_21 |
| 946 | 3'UTR | AD | DM1 | 8,173 | female |  |  | WGSrpt_13 |
| 947 | 3'UTR | AD | DM1 | 13,83 | male |  |  | WGSrpt_15 |
| 948 | Intron | AR | FRDA | 320,320 | male |  |  | WGSrpt_09 |
| 949 | Intron | AR | FRDA | 788,788 | male |  |  | WGSrpt_11 |
| 950 |  |  | (controls) |  |  | 58 | 14 | 2 |
| 951 |  |  |  |  |  |  |  |  |

### Sequencing Data generation

WGS data with PCR (WGS_PCR1 and WGS_PCR2) was generated by the Kinghorn Centre for Clinical Genomics, Garvan Institute of Medical Research, Sydney, Australia with HiSeq X Ten. The WES data (WES_PCR) was generated by the Australian Genome Research Facility, Melbourne, Australia, and sequenced on a HiSeq 2500 sequencer. All WGS_PF samples were sequenced on the Illumina HiSeq X sequencing platform at Illumina, La Jolla, California, USA. Further details can be found in Dolzhenko et al. All sequencing data was aligned to the hg19 human genome reference using the Bowtie 2 aligner^21^ in local alignment mode.

### Definition of Repeat Expansion Loci

Table 1 defines the chromosomal location, physical map location, disease, genetic disease model and repeat motif, normal and repeat expansion size for 24 repeat expansion loci, which cause neurological disorders. For the analyses in this paper we examined 21 of these STR loci, excluding the more recently discovered SCA37 and FAME1 loci, and the SCA31 locus, where the inserted repeat is not in the reference sequence. This focused the analysis on currently most likely tested expansion loci and in particular concentrating on the spinocerebellar ataxia repeat expansion loci.

### Data extraction for repeat expansions

We developed a two-step analysis method, called exSTRa, detailed in the Supplemental data, to identify individuals likely to have a repeat expansion at a particular STR locus. The analysis method extracts STR repeat content information for each read, stemming from a particular individual, which has been identified as mapping to one of the 21 STR loci. We designed a statistical test that captures the differences between an individual to be tested within a cohort of cases and controls. All N individuals within a cohort are examined in turn at each of the 21 known pathogenic repeat expansion loci by comparing each individual in turn to all N other individuals in each cohort. This generates 21xN test statistics per cohort. The empirical p-value of the test statistic was determined using a simulation method. All p-values over all STR loci for all individuals within each cohort were assessed for approximate uniform distribution with histograms and Quantile-Quantile (Q-Q) plots.

Raw data was visualized using empirical cumulative distribution functions (ECDFs), which display the distribution of the amount of STR repeat motif found in each read, ordered from smallest to largest content amount, as a step function. This allows comparison of the distributions, regardless of sequencing depth. Reads generated from expanded alleles have increased numbers of repeat motifs in their reads compared to reads stemming from normal alleles. This produces a shift of the read repeat motif distribution to the right for the individual with the repeat expansion, in comparison to reads from individuals with normal alleles.

### Simulation Study

We conducted a simulation study using the next generation sequencing data simulation package ART^22^, which simulates NGS data with realistic error profiles based on supplied reference genomes. Alleles at STR loci were simulated using reference genomes where alleles (normal, intermediate, expanded) had been inserted into the reference genome. STR loci such as HD do not have an intermediate range or only a very narrow range. We extensively searched the literature to determine pathogenic and non-pathogenic ranges of STR length alleles. We only used the ‘overall’ distribution, ignoring any ethnic specificity for these loci. We did not apply a stutter model in the simulations, as this was not feasible due to ARTs constraints. We simulated data for 20 STR loci (excluding FAME1, SCA31, SCA37 and SCA31), for 200 controls, and ten normal, ten intermediate range and ten expanded individuals. These 30 individuals were tested for expansions. The STR genotype for the controls was randomly chosen based on the distributions of these as described in the literature (Supplemental Table S3). Ten normal, intermediate and expanded alleles were chosen based on uniform distances between alleles, covering the known normal, intermediate and expanded allele ranges as described in the literature (Supplementary Table S4); for autosomal dominant loci, the second allele was chosen randomly with the same method as the controls. For the recessive STR loci EPM1 and FRDA we sampled two expanded alleles for individuals with disease. To allow for STR loci assessment on the X chromosome (FRAXA, FRAXE and SBMA) we generated half of the samples as male and the other half as female, with males having a single X chromosome and hence a single STR allele. For the X chromosome STR loci we only investigated the male individuals. To investigate the effect of control sample size on detection with exSTRa we sub-sampled the control cohort at intervals of 50, with control cohort sizes ranging from 50 to 200 individuals. The ART command used to generate the simulated data was:

art_illumina -i ${file} -p -na -l 150 -f 50 -m 450 -s 50 -o $outfile/$base −1 ${profiles}/HiSeqXPCRfreeL150R1.txt −2 ${profiles}/HiSeqXPCRfreeL150R2.txt

### Performance evaluation

For exSTRa we called individuals as being normal or expanded based on the Bonferroni multiple testing corrected p-values derived from our empirical p-values. The number of Bonferroni corrections for the four cohorts was performed based on the 21 STRs tested per individual for the WGS cohorts and 13 for the WES cohort. Repeat expansion calls were compared to the known disease status. Performance of all four methods was evaluated by examining the number of true positives (TP), true negatives (TN), false positives (FP), false negatives (FN), sensitivity, which is defined as TP/(TP+FN) and specificity, which is defined as TN/(TN+FP) at each STR and then summarized across the STR loci, within cohorts.

### Comparison with ExpansionHunter, STRetch and TREDPARSE

ExpansionHunter^17^ estimates the repeat size using a parametric model but does not attempt to call repeat expansions in a probabilistic framework. ExpansionHunter was used to determine if alleles were larger than currently known smallest disease-causing repeat expansion alleles. STRetch was used to detect the presence of repeat expansion using its statistical test, which is also an outlier detection test. Bonferroni corrections were calculated as per the exSTRa analysis. TREDPARSE was used to both estimate the repeat size and to detect the presence of an expansion based on its likelihood model. Bonferroni corrections were applied in the same way as for exSTRa.

## Results

### Simulation Study Results

The simulation study of the 20 STR loci provide evidence of the validity and robustness of the exSTRa test statistic with respect to control cohort size, repeat expansion size and known expansion status. Decreasing the control cohort in exSTRa showed that results were robust as the control sample size decreased (Supplementary Figure S5). exSTRa also showed consistent results when the size of the repeat expansion allele varied, with longer expansion alleles achieving smaller p-values (Supplementary Figure S4). Overall exSTRa p-values showed adequate Type 1 error, and good discriminatory ability between expansion and non-expansion individuals (Supplementary Figures S3 and S4). The ECDF plots, which are unique to exSTRa, show the effect of increasing expansion size in all STRs, with commensurate right shifts of the distributions. The ECDFs also allow heuristic determination of the genetic model, with larger shifts to the right for the recessive FRDA STR and the X-linked STRs (FRAXA, FRAXE and SBMA). Dominant loci only show the shift in ECDF for the upper half of the ECDF (Supplementary Figure 2). All STR loci performed well for repeat expansion detection in the simulation studies, including FRAXA and FRAXE. The simulated dataset is available to other researchers on request.

### Coverage and Alignment Results for study cohorts

Full coverage and alignment results are in Supplemental Table S2 for three cohorts, but not WGS_PF_3, which is described in Dolzhenko et al^17^. The median coverage achieved was 44, 66, 82 and 46.3 for cohorts WES, WGS_PCR_1, WGS_PCR_2 and WGS_PF_3 respectively, with 1^st^ and 3^rd^ quartile coverage of (37,48.25), (49.5,71), (76.5,84) and (44.9,47.9) Genome-wide sample specific coverage variability, as measured by the median IQR of the mean coverage library size corrected samples, was very similar between all three WGS_cohorts (WGS_PCR_1 median IQR = 8, WGS_PCR_2 median IQR = 5.7, WGS_PF_3 median IQR = 8.3). In contrast the WES data showed substantial variability (median IQR = 22.3).

### STR loci sequencing coverage ability

We examined the 21 STR loci for coverage in our four study cohorts. As expected WES_PCR only achieved reasonable coverage for repeat expansion detection in a subset of the STR loci. However, this included many of the known repeat expansion STRs located in coding regions (8 out of 10) (Figure 1 and Supplemental Figure S1). SCA6 (OMIM #183086, *CACNA1A*) and SCA7 (*ATXN7*) are poorly covered. Despite the use of the Agilent SureSelect V5+UTR capture platform, which incorporates UTRs we achieved no, or very low coverage, for the known repeat expansion loci located in the UTR, such as FRAXA (OMIM #300624), FRAXE (OMIM #309548) and DM1 (OMIM #160900). DM1 and SCA7 are not captured by the Agilent enrichment platform (Supplemental Table S3), however both FRAXA and FRAXE are targeted and therefore should be captured. In general, WGS data outperformed WES over all STR loci, with one exception, SCA3 (OMIM #109150), located in the coding region of *ATXN3*. The reason for this is currently unknown.

**Figure 1.**
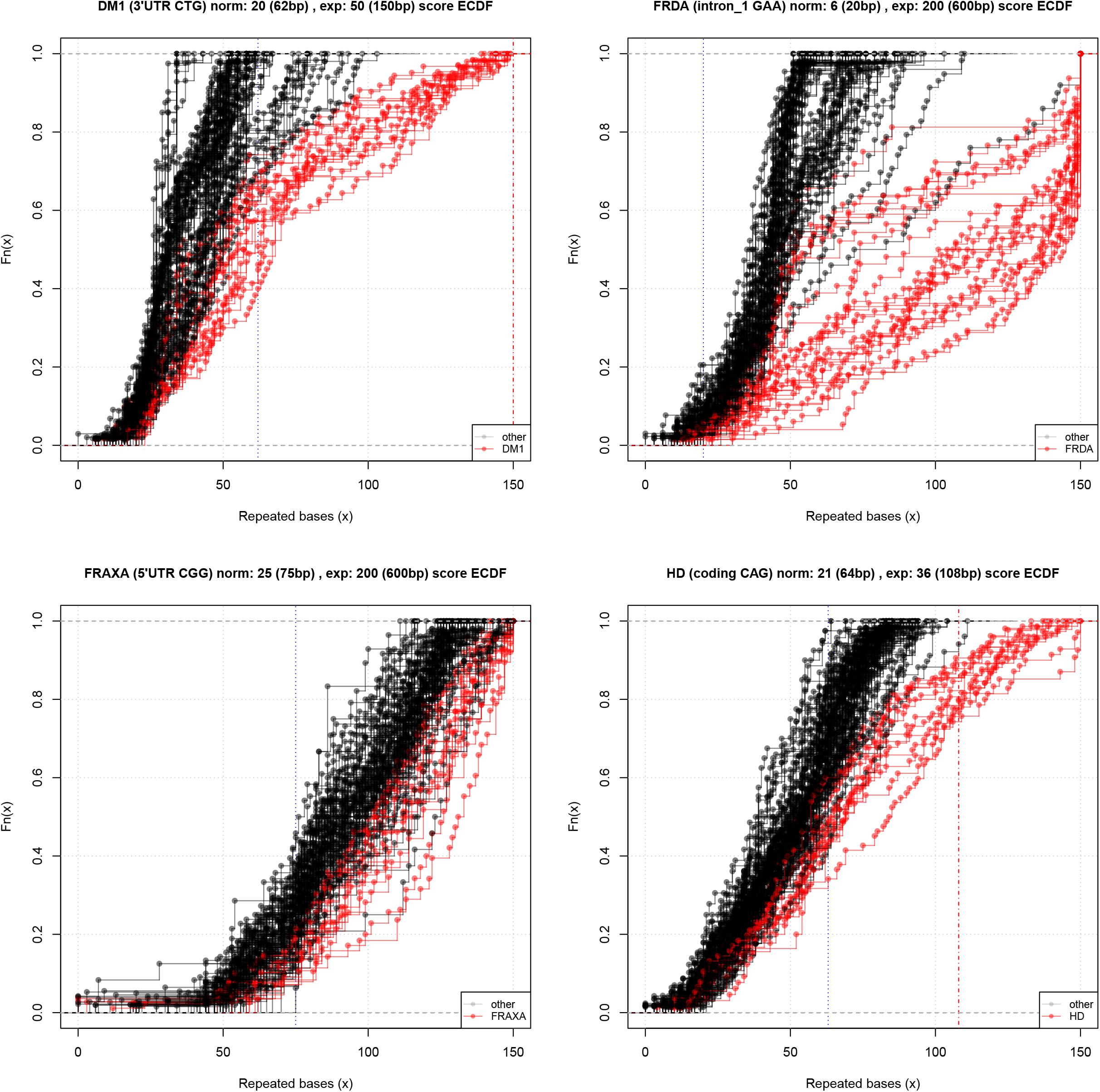
ECDF of repeat expansion composition of reads from the WES cohort, depicting four different known repeat expansion disorders captured by WES (HD, SCA2, SCA6 and SCA1). Sample rptWEHI3 (blue) is a known HD repeat expansion patient. The expanded allele size is not known. Sample rptWEHI1 (yellow) a known SCA2 repeat expansion of length 42 repeats, sample rptWEHI2 (red) a known SCA6, of length 22 repeats, and sample rptWEHI4 (green) a known SCA1 patient, of length 52 repeats. The title at the top of each individual figure gives the locus being examined, the reference number of repeats in the hg19 human genome reference with the corresponding number of bps, and the smallest reported expanded allele in the literature (with the corresponding number of bps in brackets). The blue dashed vertical line in the plot denotes the largest known normal allele, the red dashed vertical line denotes the smallest known expanded allele.

### Visualizations of repeat motif distributions

ECDF curves of selected loci are shown for each cohort to illustrate the data. Full results for all 21 loci, for all WGS cohorts, and 10 covered loci for WES cohort, are given in Supplemental Figures S6-S11. STR loci varied in their coverage with several loci consistently poorly captured. These were usually loci that are rich in GC content. Short read NGS data has a known GC bias with a GC content of 40-55% maximizing sequencing yield, depending on sequencing platform.^23^ The shape of the ECDF is affected by additional factors such as: the genetic model (dominant, recessive or X-linked) and capture efficiency (for WES).

The STR loci also showed differences in variability with regards to STR motif lengths. Some STR loci, such as SCA17 (OMIM #607136) and HDL2 (OMIM #606438), showed little variability in STR allele distributions, regardless of NGS platform in our cohorts. Identification of outliers is easier for these loci, with low background variability. Those repeat expansion disorders that are autosomal recessive or X-linked recessive (in males), also show much clearer outlier distributions (Figure 2, top right panel). This is due to the outlier distribution deviating for either both alleles, or, in the case of the X-chromosome, and only males, just the one allele having to be examined (not performed in this analysis).

**Figure 2.**
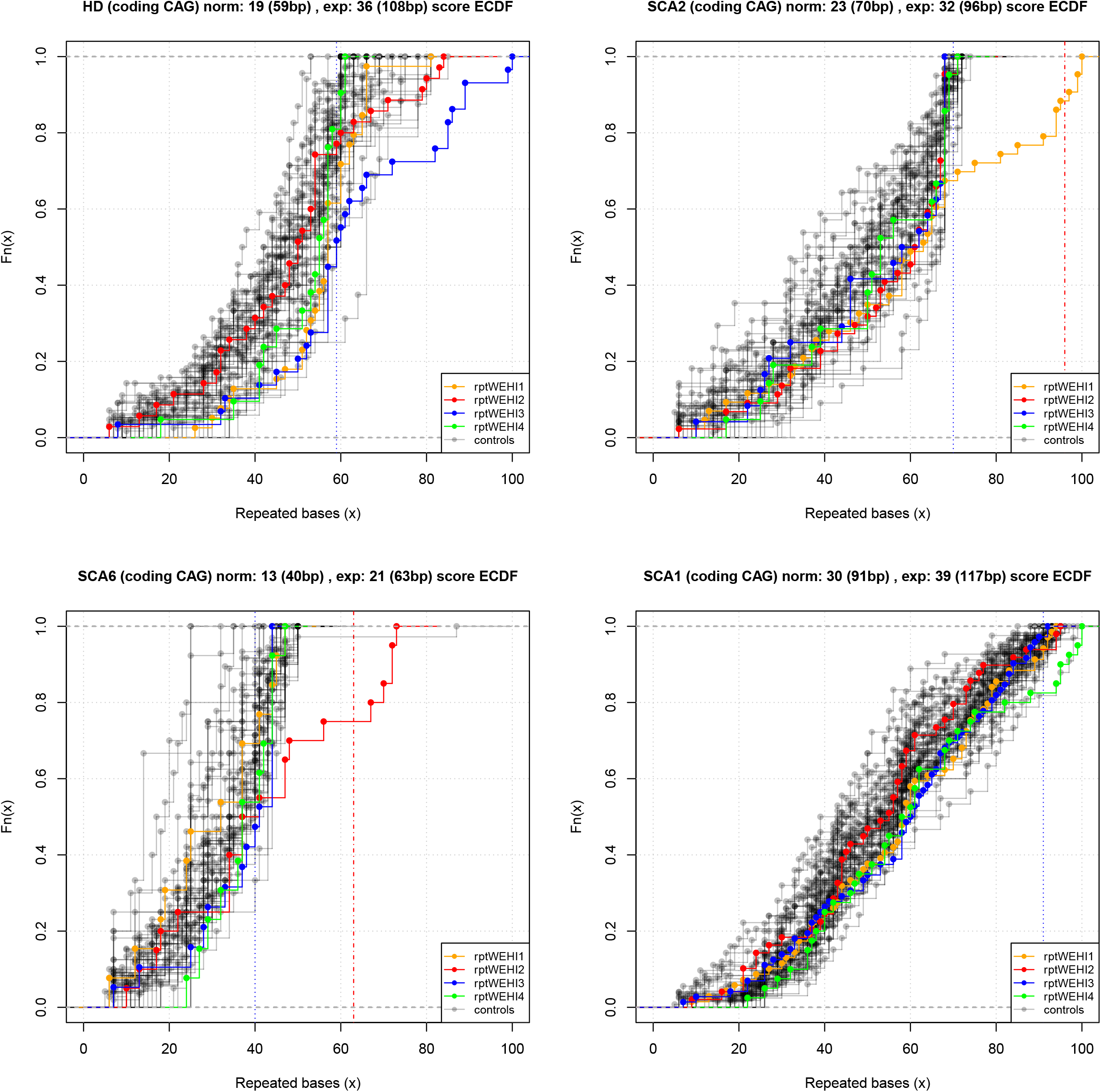
ECDFs of repeat expansion composition of reads from the WGS_PCR_2 cohort, depicting four different STR loci (top left = SCA1, length of the expanded alleles are 52 and 45 repeats; top right = FRDA, length of the expanded alleles are 320 and 788 repeats; bottom left = SCA7, length of the expanded allele is 39; bottom right = DM1, length of the expanded alleles are 173 and 83 repeats). Here coloured samples at each STR indicate those called by exSTRa as repeat expansions at the STR locus. The title at the top of each individual figure gives the locus being examined, the reference number of repeats in the hg19 human genome reference with the corresponding number of bps, and the smallest reported expanded allele in the literature (with the corresponding number of bps in brackets). The blue dashed vertical line in the plot denotes the largest known normal allele, the red dashed vertical line denotes the smallest known expanded allele.

### Statistical test results for exSTRa

Test statistics were generated for all 21 loci for all N individuals for all four cohorts with exSTRa. Combined p-values over all STR loci for all individuals within each cohort showed approximate uniform distribution with histograms (Supplemental Figure S12) and Q-Q plots (Supplemental Figure S13), albeit with some inflation of p-values at both tails. Our study cohorts had very small numbers of control individuals for some of the cohorts.

### Expansion call results

Expansion call results are presented in summary form in Tables 3 and 4, and at the individual level in Supplemental table S4 and S5. For the cohorts WES_PCR, WGS_PCR_1, WGS_PCR_2, WGS_PCR_2_30X_1, WGS_PCR_2_30X_2 and WGS_PF exSTRa achieved sensitivities of 1, 0.67, 0.81, 0.81 and 0.75 and 0.77 respectively, for these cohorts (Table 4), with very high specificity (all cohorts >0.97). Sensitivity is poorly estimated due to the small number of true positives (TPs) in some cohorts, which leads to large variability. This is particularly the case for WES_PCR (4 cases) and WGS_PCR_1 (3 cases). This has also resulted in highly variable results for the other methods. FRAXA was the STR most refractory to analysis, performing poorly regardless of sequencing platform and repeat expansion detection method. Excluding this locus in the evaluation of WGS_PF increased the sensitivity from 0.77 to 0.84, but specificity remained unchanged at 0.97.

**Table 3.**
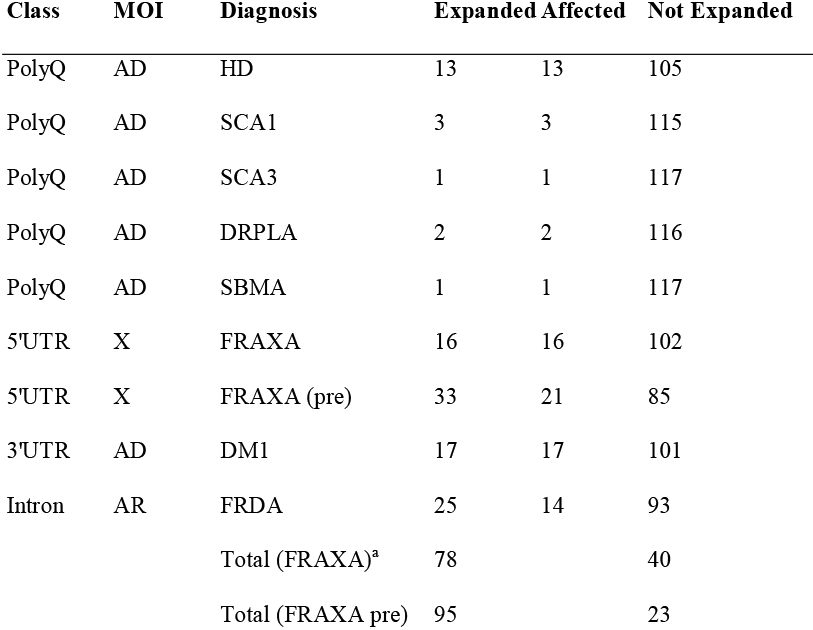
WGS_PF cohort. Cohort of 118 individuals sequenced with Illumina PCR-free library preparation. Only total number of samples are listed, rather than actual samples. Details of samples are available in Dolzhenko et al 2017. ^a^Total only includes FXS individuals, and no intermediate pre expansions.

**Table 4.** Repeat expansion detection results for all four cohorts. ^Individuals designated as controls have no known repeat expansions. Individuals designated as cases have one known repeat expansion, but are controls for all other loci tested. TP, true positive; FN, false negative, TN, true negative; FP, false positive; Sensitivity, TP/(TP+FN); Specificity, TN/(FP+TN); WES cohort labeled with (*) only assessed over ten STR loci in the capture design. WGS_PCR_2 was also analysed split into two sub-cohorts, split by flow cell lane, and are designated as WGS_PCR_2_3 0X_1 and WGS_PCR_2_3 0X_2.^a^STRetch was Bonferroni corrected for the same number of tests as the other methods, and not genome-wide corrected. ^b^TREDPARSE results are given for the repeat expansion size threshold method (TREDPARSE-T) and for the likelihood ratio test based method (TREDPARSE-L). For STR loci with recessive inheritance, samples with double expansions were designated as cases for TREDPARSE-L, which takes into account the inheritance model. ^c^For the WGS_PF cohort the original ExpansionHunter results from Dolzhenko et al were used, which make use of reads aligned with a different aligner. ^d^For the WGS_PF cohort, additional results were computed using the premutation threshold to test for FRAXA expansions with ExpansionHunter (EH FRAXA_pre), TREDPARSE-T (TP-T FRAXA_pre) and TREDPARSE-L (TP-L FRAXA_pre).

|  | Cohort | Cases | Controls^ | Method | TP | FN | TN | FP | Sensitivity | Specificity |
| --- | --- | --- | --- | --- | --- | --- | --- | --- | --- | --- |
| 971 |  |  |  |  |  |  |  |  |  |  |
| 972 | WES_PCR* | 4 | 58 | exSTRa | 4 | 0 | 607 | 9 | 1 | 0.99 |
| 973 |  |  |  | ExpansionHunter | 2 | 2 | 616 | 0 | 0.5 | 1 |
| 974 |  |  |  | STRetch<sup>a</sup> | 3 | 1 | 613 | 3 | 0.75 | 1 |
| 975 |  |  |  | TREDPARSE-T<sup>b</sup> | 4 | 0 | 585 | 31 | 1 | 0.95 |
| 976 |  |  |  | TREDPARSE-L<sup>b</sup> | 4 | 0 | 574 | 42 | 1 | 0.93 |
| 977 | WGS_PCR_1 | 3 | 14 | exSTRa | 2 | 1 | 343 | 11 | 0.67 | 0.97 |
| 978 |  |  |  | ExpansionHunter | 3 | 0 | 354 | 0 | 1 | 1 |
| 979 |  |  |  | STRetch<sup>a</sup> | 1 | 2 | 336 | 1 | 0.33 | 1 |
| 980 |  |  |  | TREDPARSE-T<sup>b</sup> | 3 | 0 | 354 | 0 | 1 | 1 |
| 981 |  |  |  | TREDPARSE-L<sup>b</sup> | 3 | 0 | 354 | 0 | 1 | 1 |
| 982 | WGS_PCR_2 | 16 | 2 | exSTRa | 13 | 3 | 352 | 10 | 0.81 | 0.97 |
| 983 |  |  |  | ExpansionHunter | 8 | 8 | 362 | 0 | 0.5 | 1 |
| 984 |  |  |  | STRetch<sup>a</sup> | 11 | 5 | 338 | 6 | 0.69 | 0.98 |
| 985 |  |  |  | TREDPARSE-T<sup>b</sup> | 12 | 4 | 362 | 0 | 0.75 | 1 |
| 986 |  |  |  | TREDPARSE-L<sup>b</sup> | 11 | 5 | 362 | 0 | 0.69 | 1 |
| 987 | WGS_PCR_2_30X_1 | 16 | 2 | exSTRa | 13 | 3 | 357 | 5 | 0.81 | 0.99 |
| 988 |  |  |  | ExpansionHunter | 8 | 8 | 362 | 0 | 0.5 | 1 |
| 989 |  |  |  | STRetch<sup>a</sup> | 11 | 5 | 340 | 4 | 0.69 | 0.99 |
| 990 |  |  |  | TREDPARSE-T<sup>b</sup> | 13 | 3 | 362 | 0 | 0.81 | 1 |
| 991 |  |  |  | TREDPARSE-L<sup>b</sup> | 9 | 7 | 362 | 0 | 0.56 | 1 |
| 992 | WGS_PCR_2_30X_2 | 16 | 2 | exSTRa | 12 | 4 | 354 | 8 | 0.75 | 0.98 |
| 993 |  |  |  | ExpansionHunter | 8 | 8 | 362 | 0 | 0.5 | 1 |
| 994 |  |  |  | STRetch<sup>a</sup> | 11 | 5 | 336 | 8 | 0.69 | 0.98 |
| 995 |  |  |  | TREDPARSE-T<sup>b</sup> | 13 | 3 | 362 | 0 | 0.81 | 1 |
| 996 |  |  |  | TREDPARSE-L<sup>b</sup> | 10 | 6 | 362 | 0 | 0.62 | 1 |
| 997 | WGS_PF | 77 | 41 | exSTRa | 60 | 17 | 2330 | 71 | 0.78 | 0.97 |
| 998 |  | 77 | 41 | ExpansionHunter<sup>c</sup> | 62 | 15 | 2395 | 6 | 0.81 | 1 |
| 999 |  | 96 | 22 | EH FRAXA_pre<sup>d</sup> | 95 | 1 | 2374 | 8 | 0.99 | 1 |
| 1000 |  | 96 | 22 | STRetch<sup>a</sup> | 62 | 15 | 2207 | 76 | 0.81 | 0.97 |
| 1001 |  | 96 | 22 | TREDPARSE-T<sup>b</sup> | 52 | 25 | 2384 | 17 | 0.68 | 0.99 |
| 1002 |  | 96 | 22 | TP-T FRAXA_pre<sup>d</sup> | 72 | 24 | 2364 | 18 | 0.75 | 0.99 |
| 1003 |  | 66 | 52 | TREDPARSE-L<sup>b</sup> | 34 | 32 | 2396 | 16 | 0.52 | 0.99 |
| 1004 |  | 72 | 46 | TP-L FRAXA_pre<sup>d</sup> | 48 | 24 | 2383 | 23 | 0.67 | 0.99 |
| 1005 | WGS_PF (no FRAXA) | 62 | 56 | exSTRa | 52 | 10 | 2231 | 67 | 0.84 | 0.97 |
| 1006 |  |  |  | ExpansionHunter<sup>c</sup> | 61 | 1 | 2292 | 6 | 0.98 | 1 |
| 1007 |  |  |  | STRetch<sup>a</sup> | 62 | 0 | 2104 | 76 | 1 | 0.97 |
| 1008 |  |  |  | TREDPARSE-T<sup>b</sup> | 52 | 10 | 2281 | 17 | 0.84 | 0.99 |
| 1009 |  | 51 | 67 | TREDPARSE-L<sup>b</sup> | 34 | 17 | 2293 | 16 | 0.67 | 0.99 |

We divided the WGS_PCR_2 cohort data into two sub-cohorts, where each sample’s data comes from a single flow cell lane that has ~30X coverage. This allowed an investigation of reproducibility, and assessment at the more standard 30X coverage. Results were highly reproducible between the two 30X replicates, with only one sample generating an alternative call between the two sequencing runs. We also observed very little change in performance between the 60X and 30X data with virtually identical sensitivity and specificity (Table 4).

### Comparison with other repeat expansion detection methods

Across all cohorts (WES_1, WGS_PCR_1, WGS_PCR_2, WGS_PF) exSTRa called the most expansions (79 out of 100 known expansions) compared to ExpansionHunter 75 expansions, STRetch 77 expansions, TREDPARSE-L 52 expansions and TREDPARSE-T 71 expansions, albeit with slightly different results in the REs identified. Excluding FRAXA exSTRa called 71 out of 82 (87%) expansions, ExpansionHunter 74 expansions, STRetch 77 expansions, TREDPARSE-L 51 expansions and TREDPARSE-T 71 expansions each. Notably, exSTRa was able to identify expanded repeats at all eleven STR expansions examined. STRetch was unable to identify the SCA6 expansions in any cohort (N=2 in WGS_PCR_2, N=1 in WGS_PCR_1 and N=1 in WES_1). SCA6 is the shortest of all known repeat expansions. These shorter expansions fail to map preferentially to the decoy chromosome for the most part, leading to the inability to call this locus. This will also apply to other short repeat expansion alleles. However the other methods found most of the SCA6 expansions, regardless of sequencing platform. All four methods performed poorly when analyzing samples with an *FMR1* expansion (FRAXA). In the WGS_PCR_1 and 2 cohorts this is due to poor coverage at the FRAXA and FRAXE loci caused by GC bias issues (Supplemental Figure S1). Although there was a clear right shift of the exSTRa ECDF plots of both the full mutation and premutation *FMR1* samples (Figure 3 bottom left panel), this was not always statistically significant. The other methods similarly performed poorly with this expansion, often failing to detect it. However, ExpansionHunter and TREDPARSE-T and -R identified pre-mutation alleles for this locus ~75% of the time. exSTRa identified 5/15 FRAXA expansions, STRetch identified none and called three of these as SCA3 expansions instead. STRetch performed equal best with ExpansionHunter in the WGS_PF cohort but was the best performer once FRAXA was ignored, finding all remaining repeat expansions, albeit with the highest false positive rate. TREDPARSE and STRetch both perform particularly well for large expansions where their use of “in-repeat reads”^17; 20^, or reads that map entirely to the repeat, is highly advantageous. exSTRa does not use this information and ExpansionHunter only uses it optionally, for large repeats. Remarkably all four methods call all 13 HD expansions correctly in the WGS_PF_3 cohort (Supplementary Table S5), suggesting highly robust detection of HD expansions for WGS data. The four methods also unanimously identify the SBMA expansion and the two DRPLA expansions.

**Figure 3.**
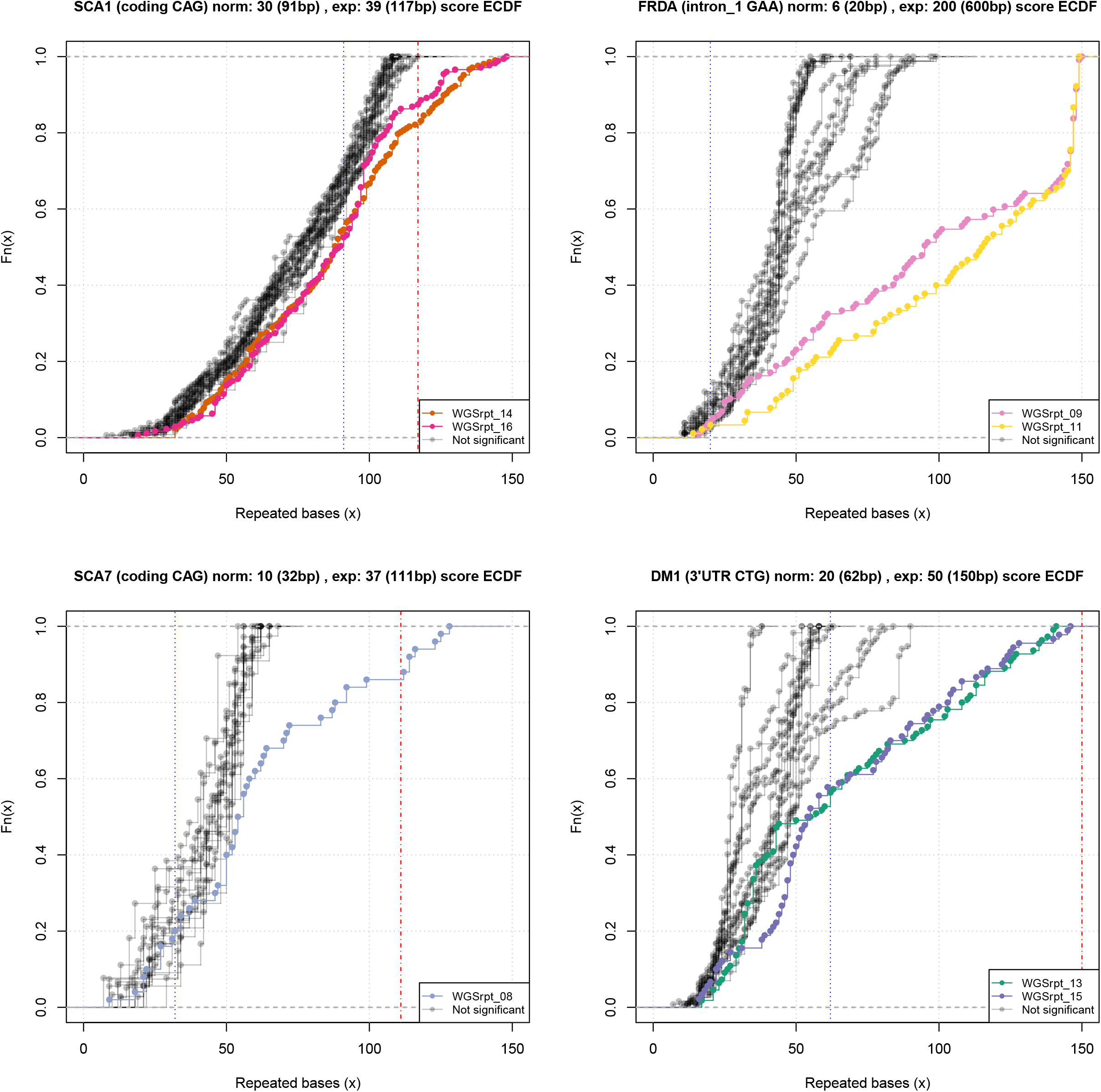
ECDFs for four repeat expansion loci from WGS_PF_3 cohort .Top left, DM1; top right, FRDA; bottom left, FRAXA; bottom right, HD .The title at the top of each individual figure gives the locus being examined, the reference number of repeats in the hg19 human genome reference with the corresponding number of bps, and the smallest reported expanded allele in the literature (with the corresponding number of bps in brackets). The blue dashed vertical line in the plot denotes the largest known normal allele, the red dashed vertical line denotes the smallest known expanded allele.

exSTRa was the equal best performing method for the WGS_PCR cohorts with TREDPARSE, and performed best overall for the WES cohort. Overall all methods performed more poorly in the WES and WGS_PCR cohorts in comparison to the WGS_PF cohort. exSTRa performs well for small repeat expansions and for platforms where small read fragments have been preferentially selected (WES_PCR, WGS_PCR). Overall the results indicate that no single method is optimal over this breadth of sequencing library preparations and STR loci. These results suggest that a consensus call that makes use of all existing methods could be advantageous. Concordance with at least one other method will be useful to maximize detection of expansions, especially since specificity is high in all WGS cohorts, across all methods (≥0.97). This drops to ≥0.93 for WES data. Using a rule whereby at least two expansion calls are required, with at least two calling methods showing concordant results to calculate a consensus call, leads to sensitivities of 1 for WES_1, 1 for WGS_PCR_1, 0.81 for WGS_PCR_2 (1, if FRAXA is excluded), 0.77 for WGS_PF_3 and 0.94 for WGS_PF_3 (excluding FRAXA) (Supplementary Tables 4 and 5, last columns).

Computational expense varied between the different repeat expansion tools. Running time for the WGS_PF cohort comprising 118 samples using 8 CPUs, was approximately 0.5 hours for exSTRa with 10^4^ permutations (12.6 hours for 10^6^ permutations), 0.6 hours for ExpansionHunter, 1.6 hours for TREDPARSE, and 2,300 hours for STRetch. STRetch requires that data is realigned to its custom reference genome, which comprises the majority of computation time and also creates additional data storage requirements.

## Discussion

Genomic medicine, which uses genomic information about an individual as part of their clinical care, promises better patient outcomes and a more efficient health system through rapid diagnosis, early intervention, prevention and targeted therapy.^24; 25^ A single affordable front-line test that is able to comprehensively detect the genetic basis of human disease is the ultimate goal of diagnostics for genomic medicine and represents the logical way forward in an era of personalized medicine. Screening tests will play a major role in the implementation of preventative medicine.

Currently, the diagnostic pathway for suspected repeat expansion disorders utilizes single gene tests or small target panels, employing a condition-by-condition approach. This method is cost effective when the clinical diagnosis is straightforward. However, for some disorders, such as spinocerebellar ataxias, the ‘right’ test is not immediately obvious.^26^ Many families remain unsolved, even after extensive genetic studies encompassing both gene sequencing and expansion repeat testing.^26^ The implementation of a single NGS-based test that could identify causal point mutations, indels and expanded STRs is likely to be cost effective in this context. NGS-based tests will act as a screening tool, to identify putative expansions, which then need to be followed up with gold-standard methods such as Southern blot analysis or repeat-primed PCR. Pathogenicity will need to be determined by clinical geneticists once the precise make-up of the repeat is determined. SNVs and indels detected in NGS also have to be validated and clinically interpreted. Detecting repeat expansions using NGS-based tests would include both increased diagnostic yield and a reduction in the diagnostic odyssey for many affected individuals.

Previously described methods such as hipSTR^14^, attempt to genotype STRs, i.e. estimate the allele sizes, which renders them ineffective when the repeat size exceeds the read length of the sequencing platform. To address this shortcoming several methods have now been developed that are designed to specifically call repeat expansions. By examining performance using >100 individuals known to have repeat expansions, spanning twelve different repeat expansion disorders, we show that exSTRa, does not require PCR-free library sequencing protocols, nor even WGS, to detect repeat expansions. We show that exSTRa delivers consistent, robust results in simulation studies.

exSTRa analysis can be run in a self contained cohort of modest size (>15 individuals). It does not require any individuals that are known to be unaffected by repeat expansions because it makes use of expanded individuals as ‘controls’ for other loci by using all available data with its robust outlier detection method. exSTRa determines significance of the outlier test statistic by simulation from the cohort using a robust estimator. Hence, the default setting for exSTRa requires that not >15% of individuals in the cohort have the same repeat expansion. exSTRa has a trimming parameter which can be adjusted. Trimming too many observations leads to non-robust results. The default setting is 15%, but this can be increased up to 50% and can be assessed for performance with the ECDF plots. This was applied to the WGS_PF cohort, which had large numbers of FRAXA (56/118, 47%) and FRDA individuals (25/118, 21%). Real disease cohorts, even ascertained from patients with diseases such as spinocerebellar ataxia, which is known to be enriched for repeat expansions, are highly unlikely to reach >15% contributions from one particular repeat expansion, based on known frequencies of such expansions.

We show that exSTRa detected the most repeat expansions across all platforms and STR loci tested. It outperforms other methods at some loci, such as FRAXE, which is the highest frequency Mendelian cause of autism. exSTRa performs well in cohorts with sequencing data with more restrictions on size-fragments and greater PCR artifacts, such as WES and WGS with PCR-based library preparations. Other advantages are that it can be run with fewer requirements (no controls necessary, no size thresholds) and its graphical ECDF representation, which allows QC and fine-tuning of analysis. The exSTRa input file is easily amended to add further loci beyond the 21 investigated. These can be determined by making use of the Tandem Repeat Finder output in the UCSC genome browser. As part of the GitHub exSTRa archive we also supply an additional input file of STRs consisting of a genome wide list of STR loci that are specifically expressed in brain. This file can be amended by the user to target specific areas of the genome, such as regions identified in linkage analysis. In comparison, ExpansionHunter and TREDPARSE (for the threshold model) currently require knowledge of the pathogenic allele size, which will not be known for novel repeat expansion loci. STRetch investigates all STRs listed in its input file simultaneously and uses its novel decoy chromosome method, facilitating genome wide analysis. However this requires re-alignment to an augmented chromosome. We also found that the decoy chromosome method does not perform well with short expansions such as SCA6, since these shorter expanded alleles will preferentially find other sites in the genome, rather than the augmented genome (data not shown). exSTRa does not attempt to call allele sizes, which TREDPARSE, ExpansionHunter and STRetch infer. However, gold standard validation with repeat-primed PCR or Southern blot still needs to occur prior to return of the genetic findings, and these methods size alleles more accurately than the NGS-based methods^27^.

We have not investigated the impact of different aligners in detail, but examination of ECDFs from the same cohort but aligned with BWA and Bowtie, the two most commonly used aligners, show highly concordant results. The ability to use existing alignments is a valuable time saving step for STR expansion analysis. exSTRa’s ECDF plots inform researchers if re-alignment is necessary or not when batches from different cohorts are combined. Combining cohorts across sequencing platforms is not advisable because motif capture and hence distributions of motif sizes differ between platforms leading to batch effects.

Some expansion alleles show population heterogeneity in allele sizes, which could influence the inference of expansions with exSTRa, but will also affect other repeat expansion detection methods since they also implicitly assume homogeneity of repeat expansion distributions. One advantage of exSTRa in this context is that the ECDF method allows assessments of the results for such features. If appropriate, population heterogeneity/membership can be assessed with methods such as PLINK^28^ or PEDDY^29^, allowing the identification and removal of population outliers or stratification of cohorts. Furthermore the exSTRa ECDF method allows assessments of the results for such features.

In the context of our results, exSTRa, and the other three methods appear to have potential as a population screening tool for carrier status. For example, all the methods should be able to identify carriers for Friedreich’s ataxia, the most prevalent of the inherited ataxias, with a carrier frequency of ~1/100 with high sensitivity and specificity. More broadly, although the current version of exSTRa performed suboptimally for detection of *FMR1* expansions, we believe these limitations can be resolved with further refinements of exSTRa or similar detection methods. Fragile X syndrome (FXS) is the most common cause of inherited ID. Approximately 1/300 individuals carry a premutation allele (55-200 repeats) which causes fragile X-associated tremor ataxia syndrome and fragile X primary ovarian insufficiency^30^. Currently, newborn/carrier screening is not performed for FXS. Historically, there was no medical advantage to early detection of FXS, although recent targeted treatments have shown potential benefits.^31; 32^ There is now discussion regarding the clinical utility of screening *FMR1* for reproductive and personal healthcare.^33^

Given that the genetic basis of disease in many affected individuals currently remains unsolved, even after extensive genetic sequencing, we recommend the introduction of a protocol, such as exSTRa, into any standard sequencing analysis pipeline and that this be run both prospectively and retrospectively. This should identify missed repeat expansions in individuals that have only been tested for a subset of common repeat expansions, which is standard clinical practice, and will also expedite the diagnosis of individuals potentially suffering from a repeat expansion disorder. There are already >20 known repeat expansion loci, but more are likely awaiting discovery. In OMIM there are additional putative SCA loci, such as SCA25 (OMIM #608703, 2p21-p13), with as yet unidentified genetic causes, but which are potentially due to novel pathogenic repeat expansions.

With large cohorts and further improvements in methodology, we believe methods such as exSTRa and future developments will facilitate the discovery of novel repeat expansion loci, which, in turn, will identify the etiology of neurodegenerative disorders in more affected individuals and families. exSTRa enables fast discovery of repeat expansions in next generation sequencing discovery cohorts including retrospective cohorts consisting mainly of WES data or WGS PCR-free library preparation data. An important new challenge lies in novel repeat expansions that are *de novo*^4; 5^, and not represented in the reference set of STRs that all four methods need to stipulate at which genomic locations to test. Addressing this current limitation of all RE detection algorithms will require refinement of existing/ the development of new bioinformatics tools.

The identification of a potentially pathogenic repeat expansion using detection methods such as exSTRa, should not replace the current diagnostic, locus-specific, PCR-based tests. Firstly, these will remain gold-standard, with higher sensitivity and specificity than the sequencing-based methods, and secondly, they give much more accurate estimates of the size of the expanded allele(s), and the makeup of the repeat, including whether there are interruptions, which has prognostic implications for the age of onset, disease progression and outcome.

We anticipate that there will be further improvements to all of the current methods that identify RE in NGS data. There are clearly sources of bias that affect certain loci that are contributing to the poor performance at some of the STRs. For instance, we observed a GC bias for the repeat expansion alleles underlying FRAXA, FRAXE and FTDALS1, with far fewer reads able to capture these repeat expansions due to their extreme GC content. Notably FRAXA and FTDALS1 had substantially improved coverage with the PCR-free protocol.

Long read WGS will see further improvements in the detection of repeat expansion alleles, allowing capture of the entire expanded allele in a read fragment, but is currently not cost-effective, being almost 10 times more expensive than the prevailing Illumina HiSeq X sequencing platform. The development of methods such as exSTRa will lead to further improvements in patient care via clinical genomic sequencing. They will also facilitate the pending era of precision/preventative medicine, when screening tests will become much more prevalent. A universal single test will be cost and time effective in comparison to the array of existing tests currently required, to test for all known mutation types.

## Appendices

### Alignment

Alignment of each pair of FASTQ files was performed with Bowtie2^21^ to the hg19 human genome reference build in very sensitive local mode, with maximum insert sizes of 800 bp for WES samples and 1000 bp for WGS samples. BAM files were sorted and merged with the Novosort tool. Duplicate marking was performed with Picard. Local realignment and base score recalibration was performed with the GATK IndelAligner tool and the Base Quality Score Recalibration tool^34^ to produce input ready BAM files.

### Software

The first step of the analysis is performed with a Perl module, called Bio::STR::exSTRa, which carries out a heuristic procedure to extract repeat content. In summary, this procedure uses the data from the reference database for the 21 loci presented in Table 1 to identify all reads that map to each of the STR loci, for each individual to be examined. The number of repeat motifs contained by each read are determined by the heuristic procedure, which examines each read for the repeat units that that STR is known to contain. This allows for some mismatches due to impure repeats and sequencing errors. Additionally, this is more computationally efficient than determining the exact repeat start and end, and is more robust as determining the edge of the repeat can be difficult near the end of a read in the presence of mismatches.

### Bio::STR::exSTRa : A heuristic procedure to extract repeat units per read

For simplicity, the following description of the data and analysis methods is only for a single locus. The algorithm is repeated independently at each locus.

Read information is extracted from a database of STR locations, such as 2–6bp repeat unit features generated using the Tandem Repeats Finder ^35^, which is also available as the Simple Repeats track of UCSC Genome Browser. Information is extracted for one STR at a time, with the following algorithm repeated for each STR:

1. The method identifies ‘anchor’ reads that facilitates identifying reads within or overlapping the STR. To qualify as an anchor, the reads are required to map within 800 bp of the STR, with the anchor orientated towards the STR. An anchor may overlap the STR.
2. The anchor-mate mapping is checked. If the anchor-mate is mapped near the STR and is not overlapping or adjacent, then the read is discarded, while those reads overlapping the STR are taken forward to the next analysis step. Sometimes the read is unmapped, or mapped to another locus, which is then recovered for further interrogation in the next step.
3. Remaining anchor-mates have their sequence content matched for the presence of the repeat unit in the correct direction, allowing for the repeat to start at any base, or phase, of the repeat unit. For example, if the repeat unit is CAG, the method can also match AGC and GCA. The number of bases found to be part of the repeat unit is counted to derive a repeat-score for that read, that is designated at a given locus as xij for sample i and read j (note that the maximum defined j depends on the sample). If both ends of a read-pair overlap within an STR, both reads undergo this procedure and each end is given a score that can be resolved during the statistical analysis of the data (the implementation in this paper did not investigate resolving these further, with both ends left in the analysis if any). An example of matching (lower case) a CAG on the opposite strand, thus matching CTG at any starting base, or phase, of the motif, i.e. CTG, TGC and GCT: CGTTCAC**ctg**GATGTGAACT**ctg**TC**ctg**ATAGGTCCCC**ctgctgctgctgctgctgctgctg**Tt**gctgc**TTT**tgctgc**TGT**ctg**AAA This 87 bp sequence has 48 bp marked (bold and lower case) as part of the repeat.
4. The method filters out reads where the score is lower than expected in random nucleotide sequences. While not precisely true, the assumption applied is that the four nucleotides are uniformly distributed and independent with respect to other positions. Short motifs are more likely to appear by chance. The method filters out scores where x_ij_<lk/4^k^, where l is the read length and k is the motif length. 800 bp has been chosen to avoid discarding reads overlapping the STR, with the insert size of read pairs having median ~360 bp. Some protocols may need to analyse reads further than 800 bp. This can be adjusted when calling the Perl module.

The output of this Perl module consists of a tab-delimited file consisting of a table where each row in the table is the repeat content of any read from a particular individual that has been identified as mapping to an STR locus that was to be investigated.

Note that these data do not represent the true size of the allele that the read has captured but where the method predicts an individual with repeat expansion allele at a particular STR locus to show an excess of reads and read content mapping to that STR.

### R package exSTRa : detecting outlier distributions of repeat content in reads

Analysis methods for the second part of the analysis method are embedded in an R package, called exSTRa (expanded STR algorithm). The output data from step 1 can be loaded and the data visualized. In particular visualizations of the data are performed with empirical cumulative distribution functions, or ECDFs.

The analysis of the samples is treated as an outlier detection problem. For the N individuals in the cohort the method compares each individual in turn to all others, including itself for robustness, for all STR loci that will be tested for repeat expansions. Since more reads with greater numbers of the repeat motif will be visible in an individual with a repeat expansion at a particular locus, the data at the repeat locus being interrogated is used in a statistical test of a difference of distribution in number of repeats that are observed for a particular individual in comparison to the set of controls. Individuals with an expanded repeat demonstrate a shift in the distribution in comparison to individuals with normal size alleles comprising their genotype for the STR locus being examined. To visualize the results, the output is plotted as empirical cumulative distribution functions (ECDFs) in R.

### Statistical Test

We developed a statistical test to detect outlier samples in comparison to a background set of samples. These outlier samples are likely to be individuals harbouring repeat expansions. To apply this test the method utilizes an empirical quantile imputation procedure, implemented in the R function quantile(). This function calculates empirical quantiles for any desired probability, for example probability = 0.5 generates the median observation in a dataset, but it is also capable of generating quantiles at probability points that have not been observed, by interpolating the probability distribution function based on the empirical observations. We make use of this function to firstly generate the same number of ‘observations’ for all samples to be tested, defined as M. In general, n is defined so that it is the largest number of observations for all of the samples, but other values could also be chosen, such as the median number of observations. The R function quantile() is applied to generate this dataset which consists of N samples, with M observations/quantiles, leading to a dataset with N by M datapoints, or quantiles. This dataset is defined as Y=(y_ij_), where y_ij_ is the repeat content of the j^th^ quantile from the i^th^ individual.

The test statistic, which we call T_i_, is defined as the average of multiple t-statistics generated at each quantile j, above a preset threshold 0 ≤ h < 1, which we usually define h = 0.5.

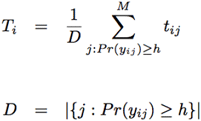

Sixteen of the 21 STR repeat expansion loci to be examined have a dominant mode of inheritance, with only one copy of the expanded allele. This can be observed with the ECDF plots for the autosomal dominant STR loci, where deviations in the repeat composition of reads are only noticeable after the median quantile, when the y-axis (which is the probability) exceeds 0.5. Observations below this threshold are likely to carry no signal, and are thus would not contribute to any test statistic attempting to discriminate between expansions and normal sized alleles.

Each quantile test statistic, t_ij_, is calculated similarly to a two-sample T-test like test statistic, but using a trimmed mean and variance, to robustly allow for the occurrence of more than one expansion in the background distribution, which is the case in the cohorts we tested but which will also likely be the case in other cohorts. The trimming percentage, or percentage of samples that are used is a parameter that can be set by the user in exSTRa, but the default is set at 0.15. Trimming is performed bilaterally, for both the lower and upper tails of the distributions, resulting in at least 30% of the samples being trimmed.

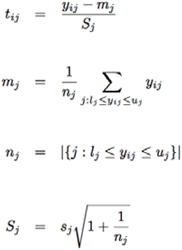

where l_i_ is the first observation included from the lower tail of the distribution after the trimmed observations and ui the last observation included from the upper tail of the distribution, with all observations beyond this trimmed. sj is the sample standard deviation of the trimmed samples.

We derive p-values for these test statistics using a simulation procedure.

Since the number of individuals in our simulations is not large and only test a single individual, standard permutation tests will not result in sufficient sampling of the empirical distribution thus resulting in a very coarse grained empirical distribution. Instead we take advantage of the well-described empirical distributions of the samples by directly simulating from the background distribution, which represents the distribution of normal, or non-expanded alleles. We perform this using robust methods to ensure that samples with expanded alleles do not influence the simulation in the simulation study.

For simulation s we simulate M quantiles for N samples, by assuming that the distributions at each quantile follow large sample theory and are thus approximately normally distributed with mean m_j_ and standard deviation d_j_, where j denotes the quantile. The method then tests this assumption by performing visual inspections of the distribution of quantiles after standardization with the R function qqnorm() and the approximation was reasonable.

The method then uses the median as our estimator for the mean, and the median absolute deviation (MAD) as our robust estimator for the standard deviation. Thus,

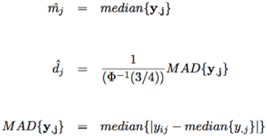

Where, and is the inverse of the cumulative distribution function of the standard normal distribution. The R function mad() incorporates the scaling factor that ensures consistency with the standard deviation when observations are normally distributed.

The method then uses the rnorm() function in R to randomly generate the N new observations for each quantile, using the STR locus and quantile specific estimators for the mean and standard deviation. The data is then sorted for each sample, as some of the new observations are no longer monotonically increasing as per definition of quantiles.

Finally, the test statistic T_s_ is calculated as defined above, but using the new data set generated from the simulation, where the first sample in the simulated data set is arbitrarily chosen to be the sample to be tested as an outlier. The method then repeat this for a desired number of simulations, say B, and then calculates the empirical p-value for our test statistic using standard methods, where:

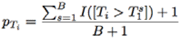

Here *I*(.) is the indicator function. T_I_^*S*^ is the test statistic for the dataset. The method calls individuals as expanded or not for each STR locus examined based on a Bonferroni corrected threshold at the 0.05 significance level, based on the number of STR tested for each sample.

Standard deviations for the empirical p-value estimator were also calculated as follows.

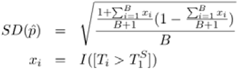

### Calling expansions with ExpansionHunter, STRetch and TREDPARSE

We performed analysis with ExpansionHunter (version 2.5.3), STRetch (GitHub commit 94d0516) and TREDPARSE (GitHub commit 83881b4), on the cohorts at the 21 repeat expansion loci listed in Table 1. The input data was the same BAM files generated as described above. Only specification files (in JSON format) for the DM1, DRPLA, FRAXA, FRDA, FTDALS1, HD, SBMA, SCA1 and SCA3 loci were provided with ExpansionHunter. The JSON files for the remaining loci were obtained by personal communication with Egor Dolzhenko (Illumina, Inc. San Diego, CA, USA). For data aligned with bowtie2, the --min-anchor-mapq parameter was set to 44, while for the original alignments of the Coriell samples this parameter was set to 60. The --read-depth parameter was set the median coverage for each sample in the WES_PCR cohort, otherwise this was computed by ExpansionHunter for the WGS samples. The list of STR loci provided with STRetch does not include FRDA, which was added manually. The EPM1 repeat motif is 12 bp and is not assessed using STRetch, which aligns to an augmented reference genome containing a decoy chromosome for each STR repeat motif up to 6 bp in size.

ExpansionHunter and TREDPARSE-T call allele lengths and genotypes. To call individuals as having expansions requires the user to define thresholds on allele sizes as to what constitutes an appropriate threshold. For FRAXA, we additionally tested using the premutation threshold (labelled FRAXA_pre), in addition to testing for full expansions. To call an expansion, we used the same thresholds as Dolzhenko et al^17^ (based on McMurray^36^) or the largest reported normal allele size at other loci. Other thresholds will change the sensitivity and specificity. TREDPARSE-L expansions calls were recorded for all samples labelled as “risk”. exSTRa p-values were Bonferroni corrected over the number of STRs tested. STRetch reports p-adjusted for multiple testing over all STRs genome wide, however unadjusted p-values were extracted and Bonferroni corrected over just the number of STRs tested. A threshold of p < 0.05 was used for significance.

## Supplemental Data

Supplemental Data includes 13 figures and 5 tables.

## Acknowledgements

We would like to thank Egor Dolzhenko and Michael Eberle (Illumina Inc), who produced the STR specification files for ExpansionHunter and gave access to the EGA00001003562. We thank Leslie Burnett, Ben Lundie, Katie Ayres and Andrew Sinclair for access to control datasets. We thank Kate Pope and Greta Gillies for assistance with recruitment and sample preparation.

RT was supported by an Australian Postgraduate Award and funding from the Edith Moffat fund. PJL was supported by NHMRC CDA2 (GNT1032364). MB was supported by NHMRC Program Grant (GNT1054618) and NHMRC Senior Research Fellowship (APP1102971). This work was supported by the Victorian Government’s Operational Infrastructure Support Program and Australian Government National Health and Medical Research Council Independent Research Institute Infrastructure Support Scheme (NHMRC IRIISS).

## Web Resources

exSTRa http://github.com/bahlolab/exSTRa

ExpansionHunter https://github.com/Illumina/ExpansionHunter

TREDPARSE https://github.com/humanlongevity/tredparse

STRetch https://github.com/Oshlack/STRetch

Picard http://broadinstitute.github.io/picard/

Novosort http://www.novocraft.com/products/novosort/

OMIM https://www.omim.org

GATK IndelAligner https://software.broadinstitute.org/gatk/

Coriell https://www.coriell.org/

## Table Legends

**Table 1** Detailed STR loci information. TRF = Tandem Repeats Finder (Benson et al, 1999). TRF match and TRF indel describe the purity of the repeat. AD = autosomal dominant, X = X-linked, AR = autosomal recessive.

**Table 2** Repeat type, genetic model, diseases, sample names and which cohorts samples appear in. Allele sizes are derived from standard laboratory tests for repeat expansions. Some individuals were not tested (Not sized) or the data was not available (not recorded). MOI, mode of inheritance; AD, autosomal dominant; X, X-linked, AR; autosomal recessive. Only the total number of controls are given denoted by (controls).

**Table 3**

Repeat Expansion detection results for exSTRa, ExpansionHunter, STRetch and TREDPARSE over all four cohorts. TP, true positive; FN, false negative; FP, false positive; TN, true negative, Sensitivity, TP/(TP+FN); Specificity, TN/(FP+TN); NA, not applicable. WES cohort labeled with (*) only assessed over eleven STR loci in the capture design. WGS_PCR_2 was also analysed split into two sub-cohorts, split by flow cell lane, and are designated as WGS_PCR_2_30X_1 and WGS_PCR_2_30X_2.

